# Machine Learning based Genome-Wide Association Studies for Uncovering QTL Underlying Soybean Yield and its Components

**DOI:** 10.1101/2021.06.24.449776

**Authors:** Mohsen Yoosefzadeh-Najafabadi, Sepideh Torabi, Davoud Torkamaneh, Dan Tulpan, Istvan Rajcan, Milad Eskandari

## Abstract

Genome-wide association study (GWAS) is currently one of the important approaches for discovering quantitative trait loci (QTL) associated with traits of interest. However, insufficient statistical power is the limiting factor in current conventional GWAS methods for characterizing quantitative traits, especially in narrow genetic bases plants such as soybean. In this study, we evaluated the potential use of machine learning (ML) algorithms such as support vector machine (SVR) and random forest (RF) in GWAS, compared with two conventional methods of mixed linear models (MLM) and fixed and random model circulating probability unification (FarmCPU), for identifying QTL associated with soybean yield components. In this study, important soybean yield component traits, including the number of reproductive nodes (RNP), non-reproductive nodes (NRNP), total nodes (NP), and total pods (PP) per plant along with yield and maturity were assessed using 227 soybean genotypes evaluated across four environments. Our results indicated SVR-mediated GWAS outperformed RF, MLM and FarmCPU in discovering the most relevant QTL associated with the traits, supported by the functional annotation of candidate gene analyses. This study for the first time demonstrated the potential benefit of using sophisticated mathematical approaches such as ML algorithms in GWAS for identifying QTL suitable for genomic-based breeding programs.

## Introduction

Soybean (*Glycine max* [L.] Merr.) is known as one of the most important legume crops with substantial economic value (Rębilas *et al.*, 2020). Soybean is widely used for food, feed, fiber, biodiesel, and green manure (Temesgen and Assefa, 2020). Despite the importance of genetic improvement in soybean yield, the soybean germplasm has in general a narrow genetic basis, especially within North American germplasm, which has resulted in limited enhancement of the genetic gain, historically (Xavier and Rainey, 2020). Therefore, there is a great need for analytical breeding to explore the optimum genetic potential of soybean (Mangena, 2020; Suhre *et al.*, 2014).

Analytical breeding strategy as an alternate breeding approach requires a better understanding of the factors, or individual traits, responsible for the development, growth, and yield (Richards, 1982). This strategy considers highly correlated secondary traits with the trait of interest as the selection criteria that can make empirical selection more efficient for improving the genetic gain (Reynolds, 2001; Richards, 1982; Xavier and Rainey, 2020). The application of the analytical approaches in plant breeding programs has been limited due mainly to lack of sufficient resources, as they are time and labor-consuming (Richards, 1982; Xavier *et al.*, 2018). Therefore, breeders are restricted to evaluating secondary traits in a small number of genotypes, which results in the limitation of the knowledge in the genome-to-phenome analysis process (Kahlon *et al.*, 2011; Nico *et al.*, 2019; Robinson *et al.*, 2009).

Yield potential in soybean is mainly determined by the following yield component traits: the total number of pods, nodes, reproductive nodes, non-reproductive nodes, and pods per plant (Pedersen and Lauer, 2004; Reynolds, 2001; Xavier *et al.*, 2018; Xavier and Rainey, 2020; Yoosefzadeh-Najafabadi *et al.*, 2021b). Of these, the total number of nodes and pods play more important roles in seed yield production than other yield components (Robinson *et al.*, 2009; Yoosefzadeh-Najafabadi *et al.*, 2021b). Several studies reported a steady increase in the total number of nodes and the total number of pods in soybean cultivars from 1920 to 2010 (Kahlon *et al.*, 2011; Suhre *et al.*, 2014; Xavier and Rainey, 2020). These findings may highlight the importance and potential use of the phenotypic and genotypic information on these traits, along with yield per se, as selection criteria in cultivar development programs (Ma *et al.*, 2001).

Genetic studies of soybean yield component traits can accelerate the breeding process more accurately (Xavier and Rainey, 2020). Genome-Wide Association Studies (GWAS), as one of the common genetic approaches, can be implemented on diverse populations to detect the quantitative trait loci (QTL) associated with the soybean yield component traits (Kaler *et al.*, 2020). Associated QTL can be used for screening large soybean populations in a short time with less elaborate efforts (Xavier *et al.*, 2018). Several GWAS algorithms have been developed for genetic studies, such as mixed linear models (MLM), multiple loci linear mixed model (MLMM), and fixed and random model circulating probability unification (FarmCPU) (Kaler *et al.*, 2020). However, due to the narrow genetic base of some plant species, including soybean, the conventional approaches may not have enough statistical power to detect reliable QTL (Kaler *et al.*, 2020; Mohammadi *et al.*, 2020; Xavier and Rainey, 2020). Therefore, the development of more sophisticated statistical methods is required in order to establish effective GWAS methods for plant species with a narrow genetic base.

Current GWAS methods are based on the conventional statistical methods that are useful for studying less complex traits in plant species with broader genetic bases (Lipka *et al.*, 2015; Pasaniuc and Price, 2017). Machine learning (ML) algorithms as powerful and reliable mathematical methods can be considered as an alternative to conventional statistical methods for performing GWAS, which are efficient for studying more complex traits in plants with narrow genetic base (Xavier and Rainey, 2020). Recently, the use of ML algorithms has been reported in different areas such as plant science (Hesami *et al.*, 2020; Yoosefzadeh-Najafabadi *et al.*, 2021a), animal science (Tulpan, 2020), human science (Chen and Verghese, 2020), engineering (Kim *et al.*, 2020), and computer science (Jordan and Mitchell, 2015). The application of ML algorithms in GWAS was previously investigated in humans by Szymczak *et al.* (2009). They explained a possible use of different ML algorithms such as artificial neural networks (ANN), Bayesian network analysis (BNA), and random forests (RF) in GWAS for human disease studies (Szymczak *et al.*, 2009). One of the most common used ML algorithms is RF developed by Breiman (2001), which generates a series of trees from the independent samples for better prediction performance (Meinshausen, 2006). The latter algorithm has been widely used in plant genomics (Ogutu *et al.*, 2011), phenomics (Yoosefzadeh-Najafabadi *et al.*, 2021a), proteomics (Jamil *et al.*, 2020), and metabolomics (Sun *et al.*, 2020).

The first and only use of the RF-mediated GWAS in soybean, for detecting the genomics association in soybean yield component traits, was reported by Xavier and Rainey (2020). Support vector machine (SVM) is another common algorithm that can detect behavior and patterns of nonlinear relationships (Auria and Moro, 2008; Hesami and Jones, 2020; Su *et al.*, 2017). Theoretically, SVM should have high performance due to the use of structural risk minimization instead of the empirical risk minimization inductive principles (Belayneh *et al.*, 2014; Yoosefzadeh-Najafabadi *et al.*, 2021a). There is a significant number of reports on the successful using of SVM in prediction problems (Denton and Salleb-Aouissi, 2020; Duan *et al.*, 2005; Hesami *et al.*, 2020; Tulpan, 2020; Yoosefzadeh-Najafabadi *et al.*, 2021a). Support vector regression (SVR) is known as the regression version of SVM that commonly used for continuous dataset. There are also reports on the successful use of SVR for addressing plant prediction problems (Awad and Khanna, 2015). However, the possible use of SVR in GWAS is still unexplored in plant science area.

In this study we aimed to: (1) gain a better understanding of the genetic relationships between soybean yield and its component traits, and (2) investigate the potential use of RF and SVM algorithms in GWAS for discovering QTL underlying soybean yield components as compared to conventional GWAS methods of MLM and FarmCPU. The results of this study will help soybean breeders to have a better perspective of exploiting ML algorithms in GWAS studies, and may offer them new genomic tools for screening high yielding genotypes with improved genetic gain based on genomic regions associated with yield components.

## Materials and Methods

### Population and experimental design

An GWAS panel of 250 soybean genotypes was grown at the University of Guelph, Ridgetown Campus in two locations, Palmyra (42°25′50.1“N 81°45′06.9“W, 195 m above sea level) and Ridgetown (42°27′14.8“N 81°52′48.0“W, 200m above sea level) in Ontario, Canada, in two consecutive years, 2018 and 2019. The panel used in this study consisted of the main germplasm of the soybean breeding program at the University of Guelph, Ridgetown Campus, that has been established over 35 years for cultivar development and genetic studies. The randomized complete block design (RCBD) with two replications was used for all four environments. In general, there were 500 and 1000 research plots per environment and year, respectively. Each plot consisted of five 4.2 m long rows with 57 seeds per m^2^ seeding rate.

### Phenotyping

In this experiment, soybean seed yield (t ha^−1^ at 13% moisture) for each plot was estimated by harvesting three middle rows. Soybean seed yield components, including the total number of reproductive nodes per plant (RNP), the total number of non-reproductive nodes per plant (NRNP), the total nodes per plant (NP), and the total number of pods per plant (PP), were measured using 10 randomly selected plants from each plot. The maturity was recorded as the number of days from planting to physiological maturity (R7, (Fehr and Caviness, 1971) for each genotype.

### Genotyping

Young trifoliate leaf tissue for each soybean genotype from the first replication of the trail at the Ridgetown in 2018, were collected and in a 2 mL screw-cap tube. The leaf samples were freeze-dried for 72 hours, using the Savant ModulyoD Thermoquest (Savant Instruments, Holbrook, NY). By using the DNA Extraction Kit (SIGMA®, Saint Louis, MO), DNA was extracted for soybean genotypes, and the quantity of DNAs was checked via Qubit® 2.0 fluorometer (Invitrogen, Carlsbad, CA). For genotyping-by-sequencing (GBS), DNA samples were sent to Plate-forme D’analyses Génomiques at Université Laval (Laval, Quebec, Canada). The GWAS panel was genotyped via a GBS protocol based on the enzymatic digestion with *ApeKI* (Sonah *et al.*, 2013). Single-nucleotide polymorphisms (SNPs) were called by the Fast GBS pipeline (Torkamaneh *et al.*, 2020), using Gmax_275_v2 reference genome. Markov model was used to impute the missing loci, and SNPS with a minor allele frequency (MAF) less than 0.05 were removed below the threshold. In total, after checking the quality of reading sequence and removing SNPs with more than 50% heterozygosity, 23 genotypes were eliminated from the experiment and 17,958 high-quality SNPs from 227 soybean genotypes used for genetic analysis.

### Statistical analyses

The best linear unbiased prediction (BLUP) as one of the common linear mixed models (Goldberger, 1962) was used to estimate the genetic values of each soybean genotype. Also, R package *lme4* (Bates *et al.*, 2014) was used to analyze yield and yield components with ‘environment’ as a fixed effect and ‘genotype’ as a random effect. To control for the possible soil heterogeneity among the plots within a given block and reduce the associated experimental errors, nearest-neighbor analyses (NNA) was used as one of the common error control methods (Bowley, 1999; Katsileros *et al.*, 2015; Stroup and Mulitze, 1991). Outliers were determined in the raw dataset based on the protocols proposed by Bowley (1999) and treated the same as missing data points in the analysis. Overall, the following statistical model was used in this study:

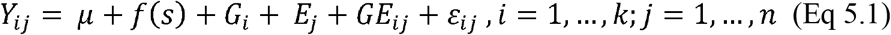

Where Y_ij_ stands for the trait of interest (soybean seed yield and yield component traits) as a function of an intercept *μ*, f(s) stands for the spatial covariate, G_i_ is the random genotype effect, E_j_ stands for the fixed environment effect, GE_ij_ is the genotype × environment interaction effect, and ε_ij_ stands for the residual effect.

The heritability was calculated for soybean seed yield and yield components using *lme4* open-source R package (Bates *et al.*, 2007) based on the following equation:

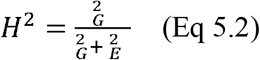

where ^2^ _G_ stands for the genotypic variance, and ^2^ _E_ is the environmental variance.

### Analysis of population structure

A total of 17,958 high-quality SNPs from 227 soybean genotypes were used to conduct population structure analysis using fastSTRUCTURE (Raj *et al.*, 2014). Five runs were conducted for K set from 1 and 15 to estimate the most appropriate number of subpopulations by using the K tool from the fastSTRUCTURE software.

### Association studies

Since different GWAS methods may capture different genomic regions (Yang *et al.*, 2018). Therefore, MLM and FarmCPU (two most common GWAS methods) and RF and SVM (two most common machine learning algorithms) were used in this study. MLM and FarmCPU were implemented by using *GAPIT* package (Lipka *et al.*, 2012), and RF, as well as SVM, were conducted through the *Caret* package (Kuhn *et al.*, 2020) in R software version 3.6.1. A brief description of each of the GWAS methods is provided below:

#### Mixed Linear Model (MLM)

This GWAS is based on the likelihood ratio between the full model, consisting of the marker of interest, and the reduced model, which is known as the model without the marker of interest (Wen *et al.*, 2018).

#### Fixed and random model circulating probability unification (FarmCPU)

This GWAS takes the advantages of using MLM as the random model, and stepwise regression as the fixed model iteratively (Liu *et al.*, 2016). False discovery rate (FDR) is used for setting the threshold both in the FarmCPU and MLM models (Benjamini and Hochberg, 1995).

#### Random Forest (RF)

This machine-learning algorithm was first implemented by Xavier and Rainey (2020) in a soybean GWAS study. This method is known as the powerful non-parametric regression approach that is derived from aggregating the bootstrapping in various decision trees (Breiman, 2001). In this experiment, a 1000-set of decision trees constructed the forest, and the GWAS analysis was done by measuring the importance of each feature (Botta *et al.*, 2014), which was an SNP in this study.

#### Support vector regression (SVR)

This machine learning algorithm is known as one of the common supervised learning methods in prediction problems (Cortes and Vapnik, 1995). This algorithm is based on constructing a set of hyperplanes that can be useful in regression problems (Fletcher, 2009). The association statistics in this algorithm can be achieved by estimating the feature importance that was previously proposed by Weston *et al.* (2001). In this experiment, SNP markers were selected as inputs, and the traits were selected as target variables for estimating the feature importance.

### Variable Importance measurement

As one of the common indices for tree-based algorithms, the impurity index was chosen as the metric of the feature importance for the RF algorithm. Regarding the SVR algorithm, the variable importance method for SVR Weston *et al.* (2001) was implemented in this dataset. For both algorithms, the importance of each SNP was scaled based on 0 to 100 percent scale. Since there is no confirmed way of defining the significant threshold in the tested algorithms, the global empirical threshold that provides the empirical distribution of the null hypothesis (Churchill and Doerge, 1994; Doerge and Churchill, 1996) was used for establishing threshold in this study. The global empirical threshold was estimated based on fitting the ML algorithm, storing the highest variable importance, repeating 1000 times, and select the SNPs based on α=0.05.

### Data-driven model processes

In order to estimate the feature importance in RF and SVR algorithms, a five-fold cross-validation strategy (Siegmann and Jarmer, 2015) with ten repetitions was applied on the dataset. All of the tested machine learning algorithms were optimized for their parameters for this dataset accordingly.

### Functional annotation of candidate SNPs

For each tested GWAS model, the flanking regions of each QTL was determined using LD decay distance (Fig.1), and then potential candidate genes were retrieved using the *G. max* cv. William 82 reference genome, gene models 2.0 in SoyBase (https://www.soybase.org). After listing potential candidate genes in defined windows around each significant SNP, at the peak of each QTL, Gene Ontology annotation, GO term enrichment (https://www.soybase.org), and the report from previous studies were used as the criteria to select and report the most relevant candidate genes associated with the identified QTL. The Electronic Fluorescent Pictograph (eFP) browser for soybean (www.bar.utoronto.ca) was also used to generate additional information such as tissue- and developmental-stage dependent expression (based on transcriptomic data from Severin *et al.* (2010)) for the identified candidate genes.

**Fig. 1.**
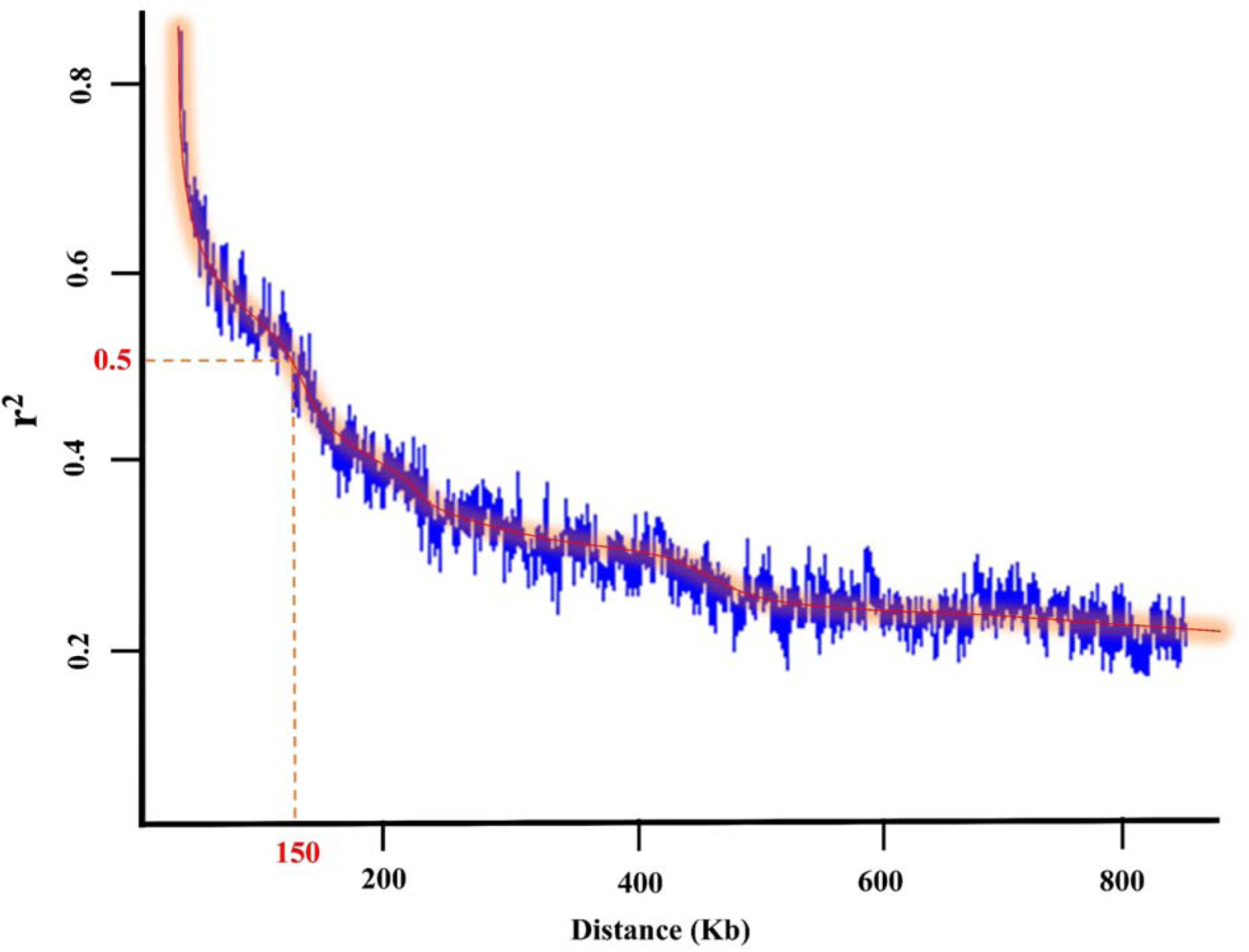
LD decay distance in the tested 227 soybean genotypes

### Visualization

All of the visualizations in this study were conducted using the *ggplot2* package (Wickham, 2011) in R version 3.6.1 software and Microsoft Excel software (2016).

## Results

### Phenotyping evaluations

The tested GWAS panel of 227 soybean genotypes showed significant variations among the genotypes for seed yield, maturity, and yield component traits. The distribution of the phenotypic measures for the traits across the four environments is presented in Fig. 2. The highest heritability was observed for maturity with an estimate of 0.78 followed by 0.34, 0.33, 0.31, and 0.30 for NP, RNP, NRNP, and PP, respectively (Fig. 2). The lowest heritability was estimated for yield with a value of 0.24 (Fig. 2). Soybean seed yield and PP showed the highest variability across the environments (Fig. 2).

**Fig. 2.**
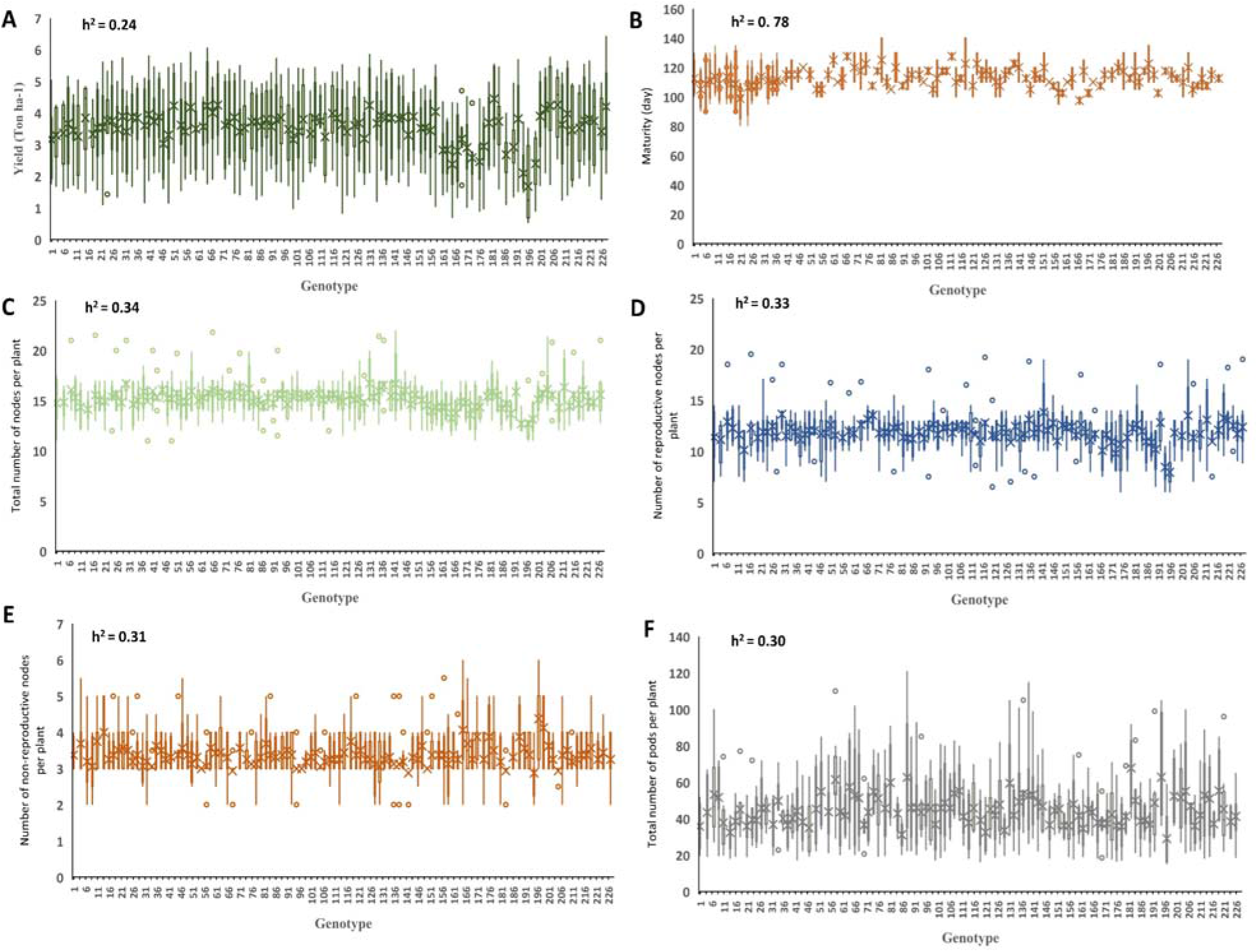
The distribution of seed yield (A), maturity (B), NP (C), NRNP (D), RNP (E), and PP (F) in 227 soybean genotypes across four environments. The estimated heritability is provided for each of the six traits. RNP: Total number of reproductive nodes per plant, NRNP: The total number of non-reproductive nodes per plant, NP: The total nodes per plant, PP: The total number of pods per plant.

The linear correlations among all the measured traits were estimated using the coefficients of correlation (*r*). Based on the results (Fig. 3), all traits were positively correlated with each other, except the NRNP that was negatively associated with yield, maturity, RNP, NP, and PP. NP showed the highest correlation with the RNP (*r*= 0.97) and NRNP (*r*= −0.63). RNP had the highest correlation (*r* =0.86) with yield among all the tested yield components (Fig. 3).

**Fig. 3.**
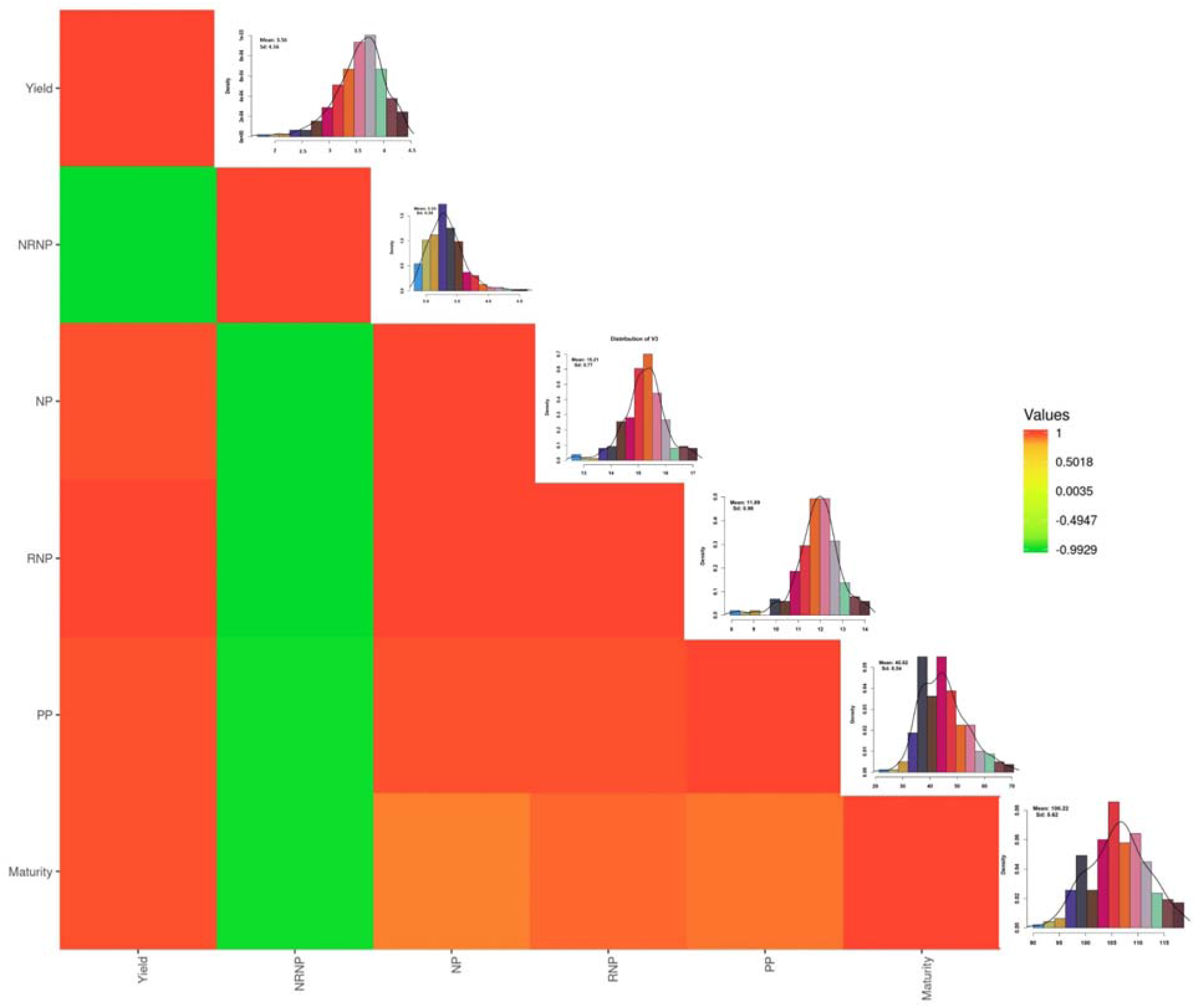
The distributions and Pearson correlations among the soybean seed yield, maturity, and yield component traits. RNP: Total number of reproductive nodes per plant, NRNP: The total number of non-reproductive nodes per plant, NP: The total nodes per plant, PP: The total number of pods per plant. The heat map scale for values is provided by colour for the panel.

### Genotyping evaluations

For the tested GWAS panel, high-quality SNPs were obtained from 210M single-end Ion Torrent reads that were proceeded with Fast-GBS.v2. From a total of 40,712 SNPs, 17,958 SNPs were polymorphic and mapped to 20 soybean chromosomes. The minimum and maximum number of SNPs were 403 and 1780 on chromosomes 11 and 18, respectively. Overall, the average number of SNPs across all the 20 chromosomes was 898, with the mean density of one SNP for every 0.12 cM across the genome.

### Population structure and kinship

The structure profile for the tested population is presented in Fig. 4. The result of genotypic evaluations suggested that the tested GWAS panel was composed of four to seven subpopulations. Therefore, we chose to conduct the structure analysis using K=7 as the appropriate K for the structure profile of the tested GWAS panel (Fig. 4). In order to reduce the confounding, the kinship was also estimated between genotypes of the GWAS panel.

**Fig. 4.**
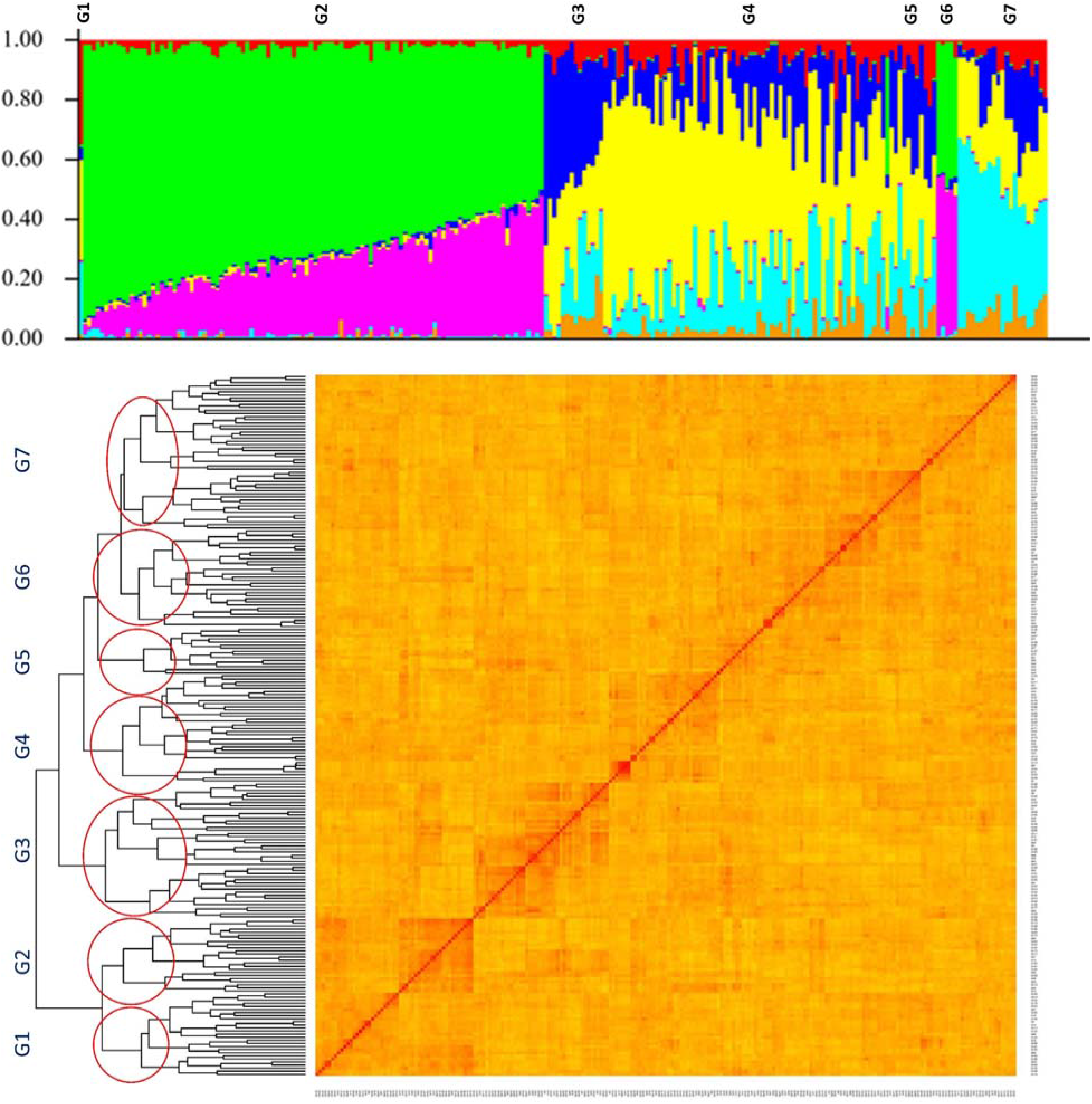
Structure and kinship plots for the 227 soybean genotypes. The x-axis is the number of genotypes used in this GWAS panel, and the y axis is the membership of each subgroup. G1-G7 stands for the subpopulation.

### GWAS analysis

The average value for soybean maturity in the tested GWAS panel was 106 days with a standard deviation of 5 days (Fig. 3). Association analysis by the MLM method identified nine associated SNP markers located on chromosomes 2 and 19 (Fig. 5A). Using FarmCPU, a total of nine associated SNP markers were located on chromosomes 2, 19, and 20 (Fig. 5A). By using the RF method, the total of three SNP markers on chromosomes 3, 7, and 16 were associated with the soybean maturity, whereas SVR-mediated GWAS detected 10 associated SNP markers located on chromosomes 2, 6, 10, 16, 19, and 20 (Fig. 5A).

**Fig. 5.**
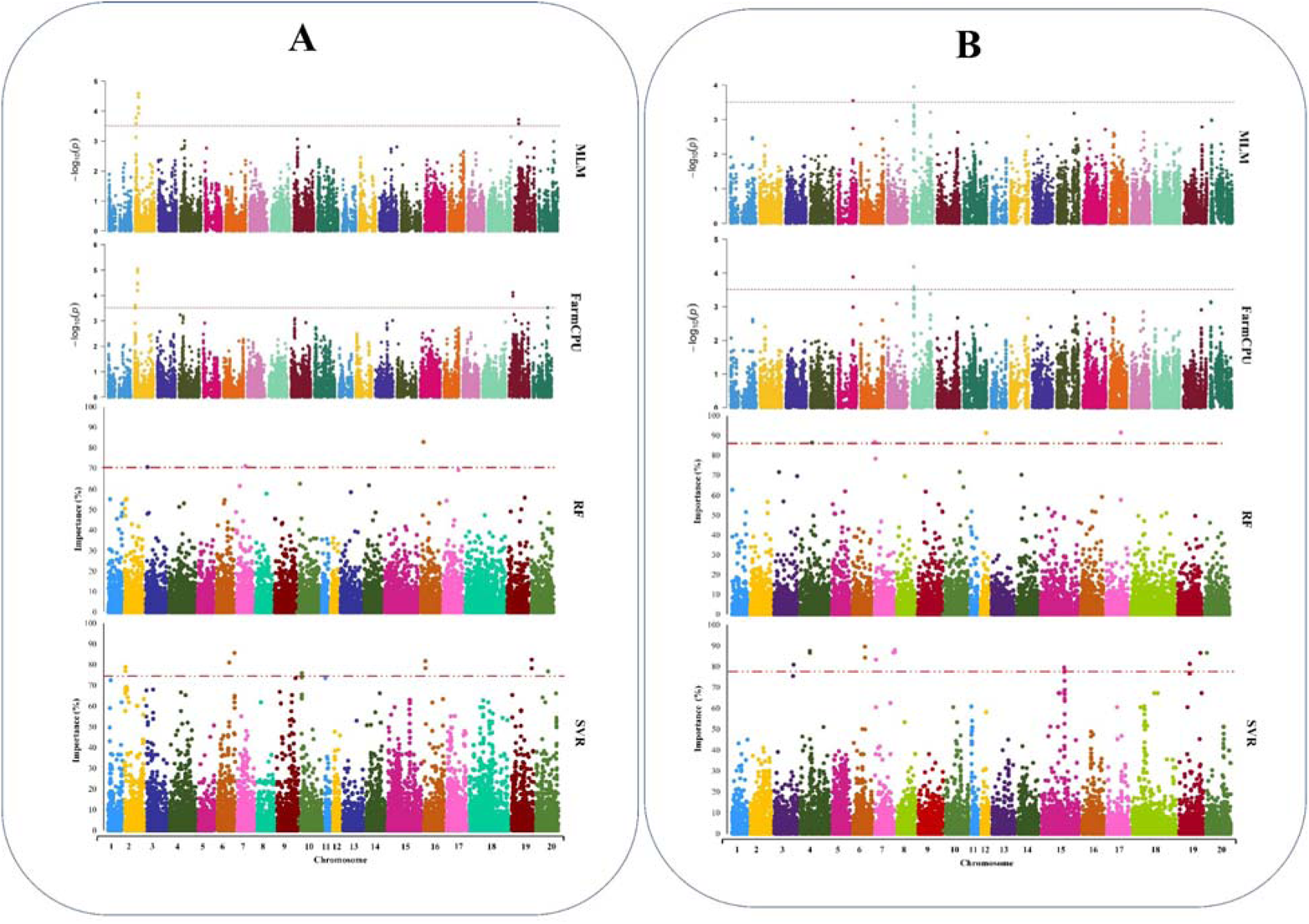
Genome-wide Manhattan plots for GWAS studies of A) maturity and B) seed yield in soybean using MLM, FarmCPU, RF, and SVR methods, from top to bottom, respectively.

SVR-mediated GWAS detected five QTL directly related to the reproductive period and R8 full maturity (Table 1). The average soybean seed yield in the GWAS panel was 3.5 t ha^−1^ with a standard deviation of 0.45 (Fig. 3). Using MLM, FarmCPU, RF, and SVR approach, we identified two, three, five, and 18 SNP markers associated with the yield, respectively (Fig. 5B). The SNP markers identified by MLM and FarmCPU were located on chromosomes 6 and 8. Using the RF-mediated GWAS method, associated SNP markers were located on chromosomes 4, 7, 12, and 17. By using the SVR-mediated GWAS method, the SNP markers were located on chromosomes 3, 4, 6, 7, 15, 19, and 20 (Fig. 5B). In SVR-mediated GWAS, the identified QTL were co-localized with eight previously reported related QTL such as seed yield, seed weight, and seed set (Table 2). However, other tested GWAS methods could not co-localized with any QTL associated with seed yield (Table 2).

**Table 1.**
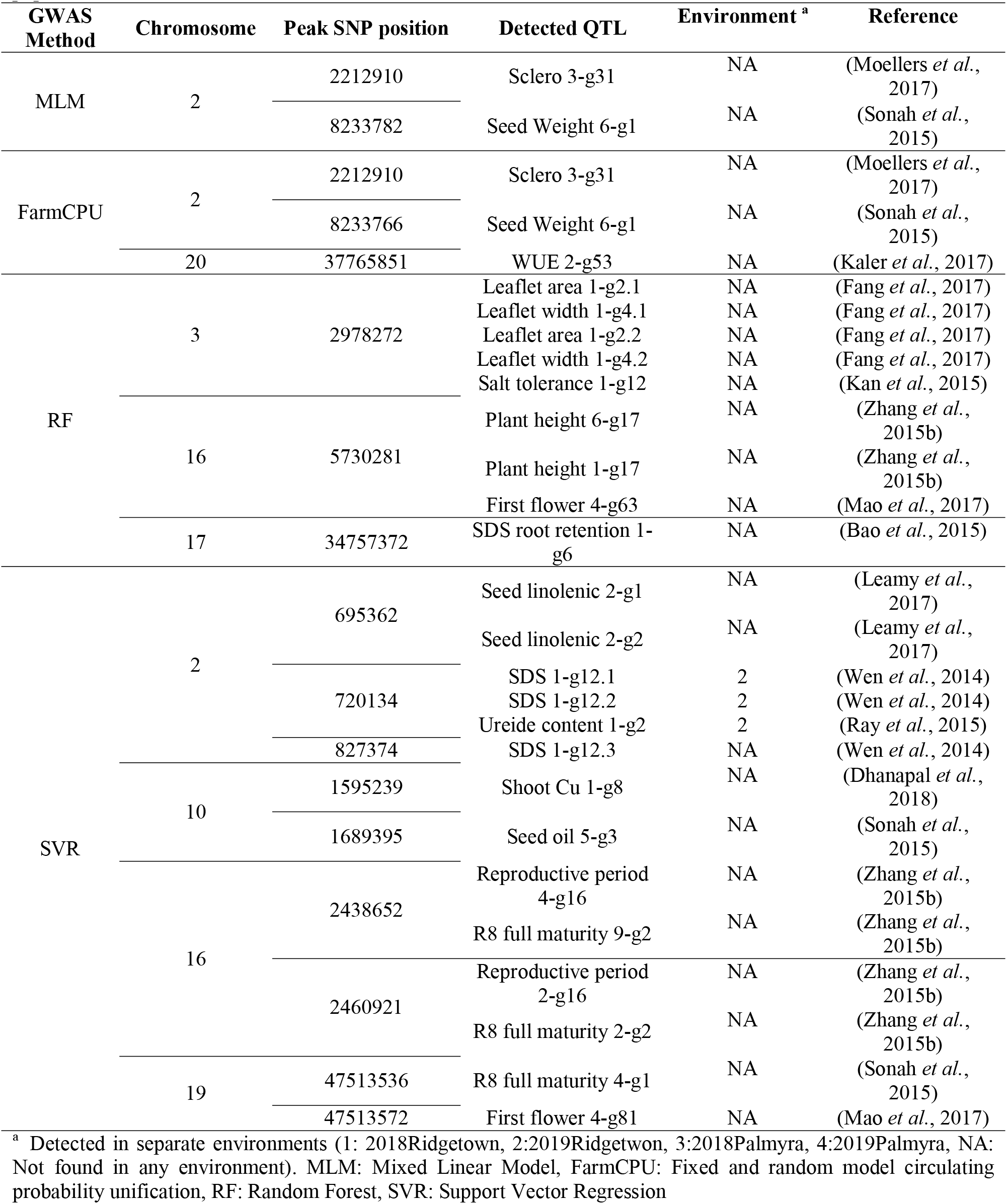
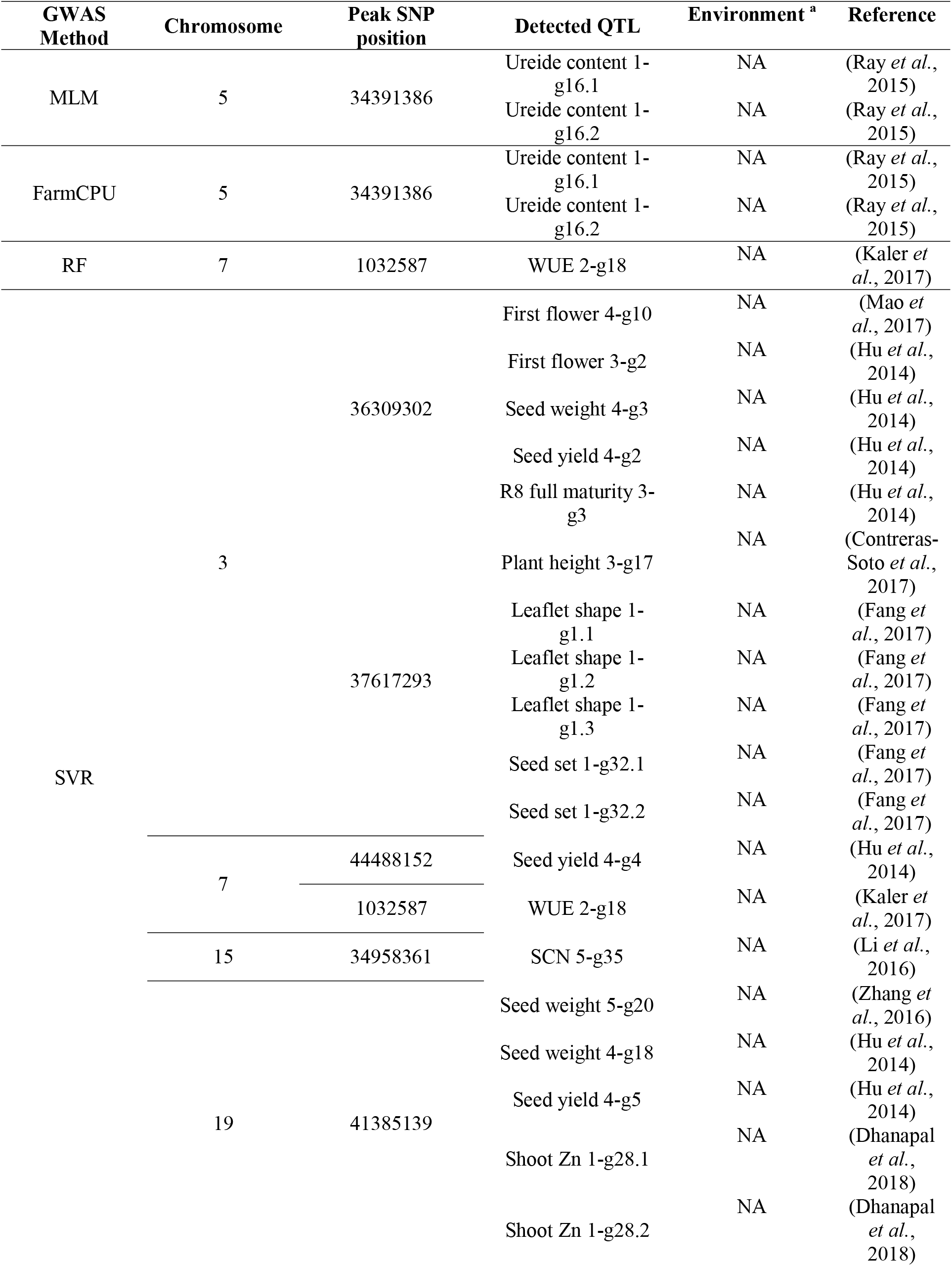
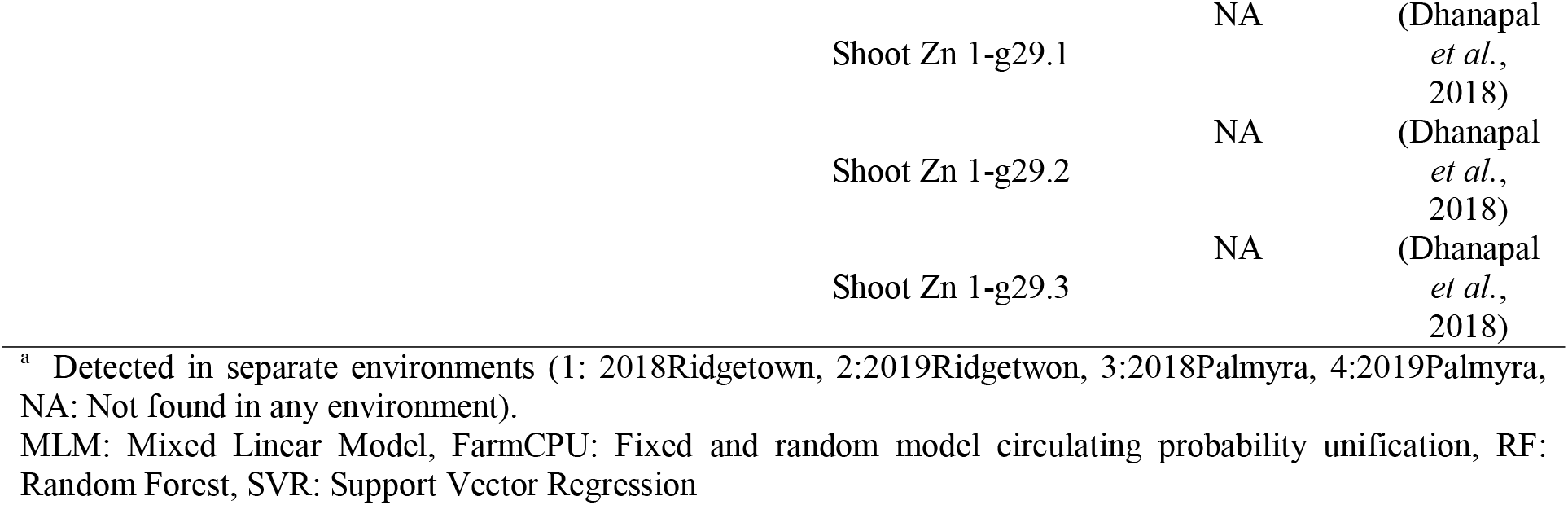
The list of detected QTL for soybean maturity using different GWAS methods in the tested soybean population.

**Table 2.**
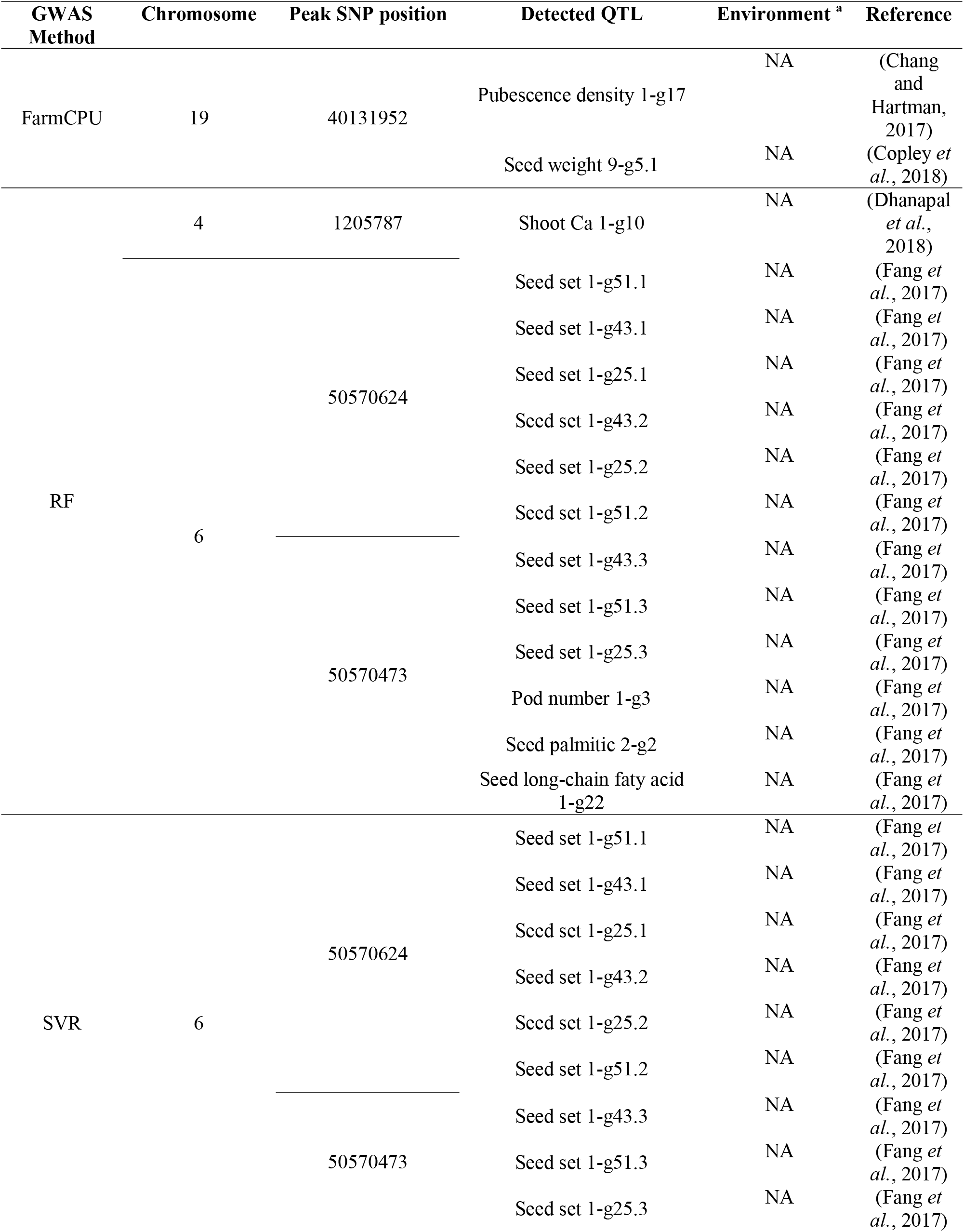
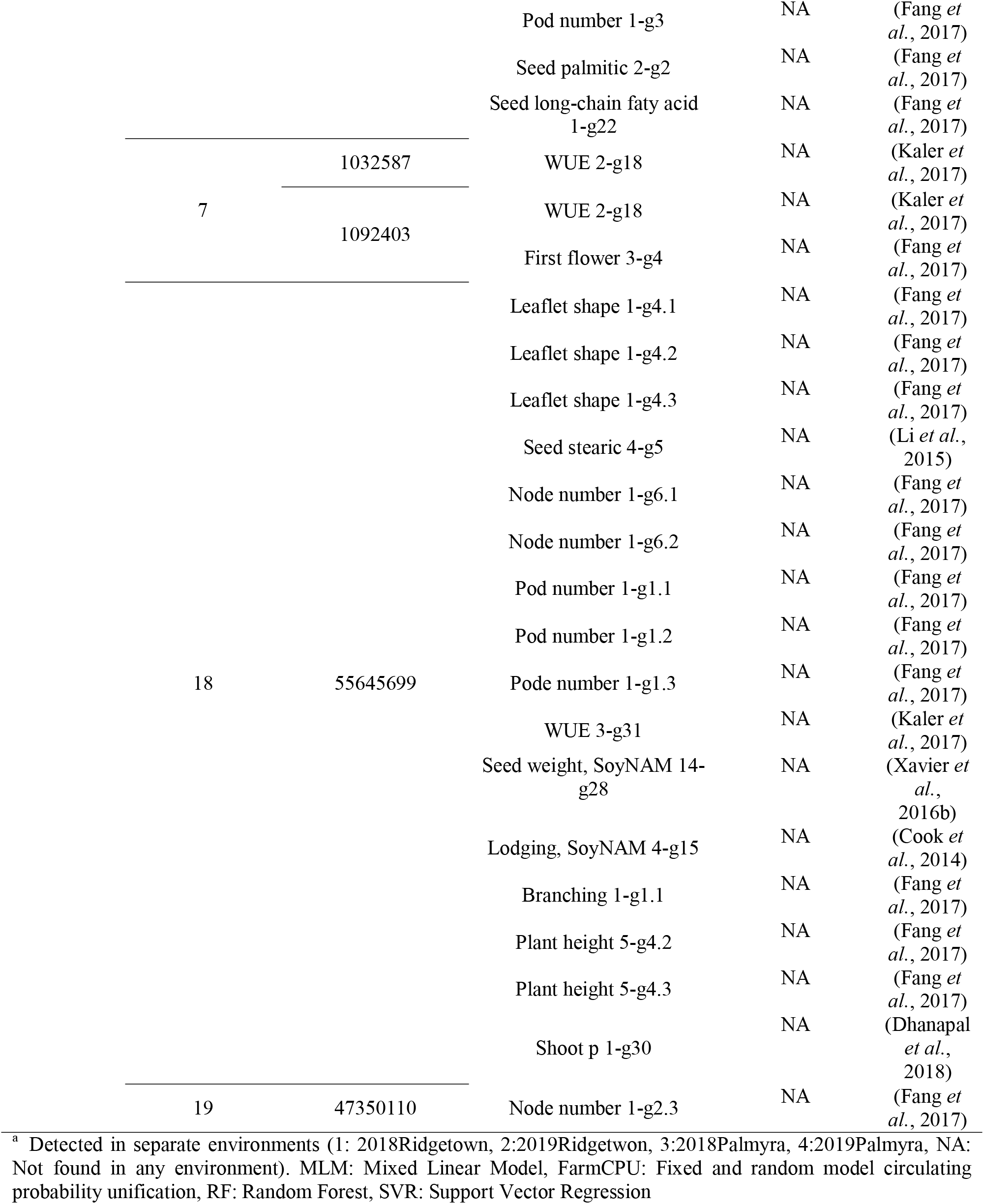
The list of detected QTL for soybean total number of nodes per plant (NP) using different GWAS methods in the tested soybean population.

The average NP in the tested GWAS panel was 15.21 nodes with a standard deviation of 0.77 nodes (Fig. 3). By using the MLM and FarmCPU methods, one and two associated SNP markers were detected, respectively (Fig. 6A). Four and ten associated SNP markers were detected by NP using RF and SVR methods, respectively. SVR-mediated GWAS was the only method that were co-localized with three previously reported NP-related QTL (Table 3). The average NRNP was 3.33 nodes with a standard deviation of 0.28 nodes (Fig. 3). A total of two, three, five, and ten associated SNP markers were detected using the MLM, FarmCPU, RF, and SVR methods, respectively (Fig. 6B). The detected SNP markers using the SVR method were located on chromosomes 4, 7, 18, 19, and 20, whereas SNP markers identified through RF were located on chromosomes 1, 4, 7, 18, and 19 (Fig. 6B). Chromosomes number 4, 8, and 15 were identified as carrying SNP markers with NRNP using FarmCPU. The MLM method identified SNP markers located on chromosomes 8 and 15, which most of the detected QTL co-localized with previously reported QTL related to seed weight, seed protein, water use efficiency, first flower, and soybean cyst nematode (Table 4).

**Fig. 6.**
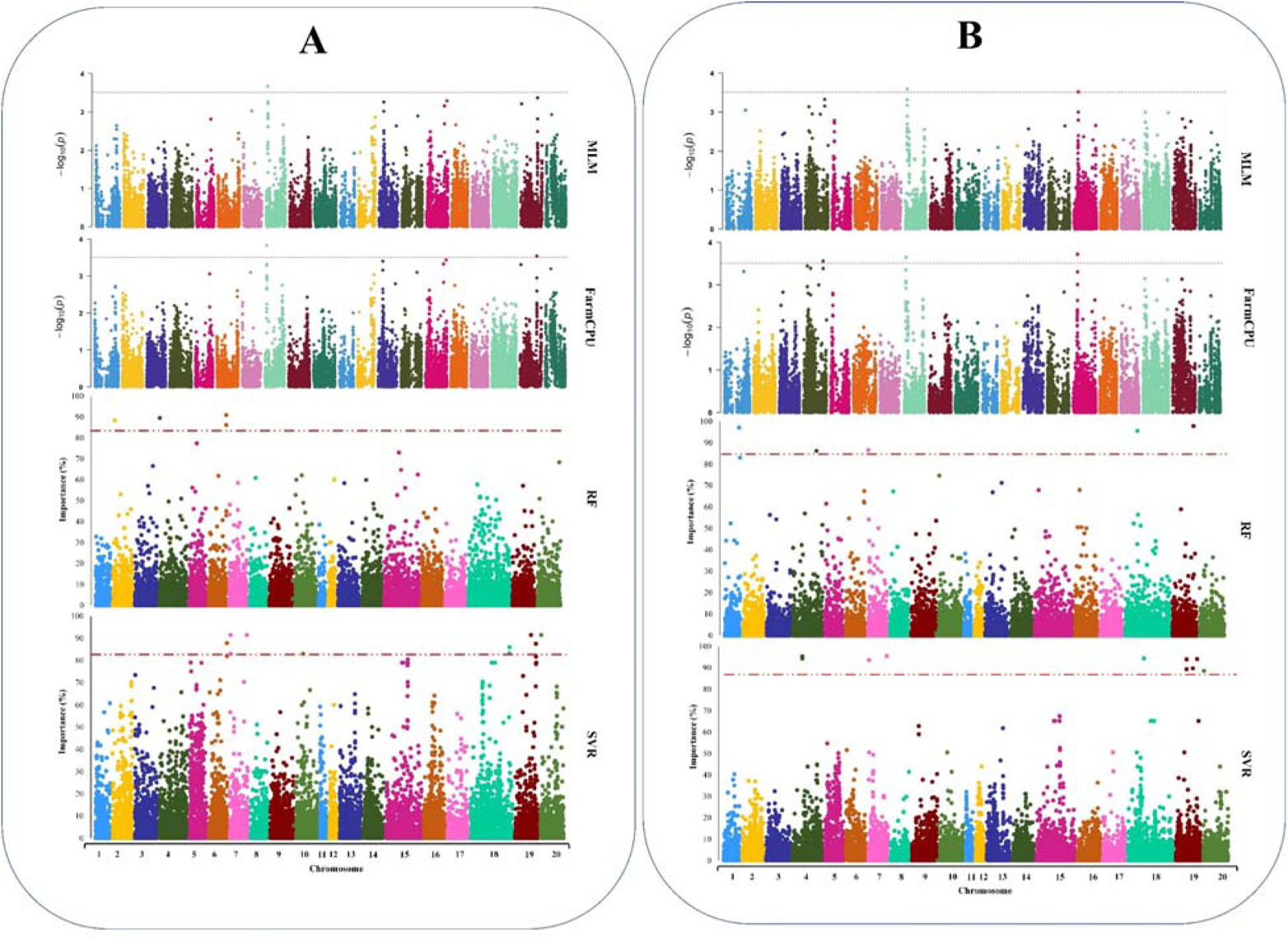
Genome-wide Manhattan plots for GWAS studies of A) the total number of nodes (NP) and B) the total number of non-reproductive nodes (NRNP) in soybean using MLM, FarmCPU, RF, and SVR methods, from top to bottom, respectively.

**Table 3.** The list of detected QTL for soybean total number of non-reproductive nodes per plant (NRNP) using different GWAS methods in the tested soybean population.

| GWAS Method | Chromosome | Peak SNP position | Detected QTL | Environment <sup>a</sup> | Reference |
| --- | --- | --- | --- | --- | --- |
| MLM | 15 | 10193796 | Seed protein 6-g2 | NA | (Zhang *et al.*, 2018) |
|  |  |  | Seed Arg 1-g4 | NA | (Zhang *et al.*, 2018) |
|  |  |  | Seed coat luster 1-g1.3 | NA | (Fang *et al.*, 2017) |
| FarmCPU | 15 | 10193796 | Seed protein 6-g2 | NA | (Zhang *et al.*, 2018) |
|  |  |  | Seed Arg 1-g4 | NA | (Zhang *et al.*, 2018) |
|  |  |  | Seed coat luster 1-g1.3 | NA | (Fang *et al.*, 2017) |
| RF | 1 | 54647498 | First flower 4-g2 | NA | (Mao *et al.*, 2017) |
|  | 7 | 329800 | Phytoph 2-g32 | NA | (Qin *et al.*, 2017) |
|  |  |  | Phytoph 2-g7 | NA | (Qin *et al.*, 2017) |
|  | 18 | 12945778 | SCN 4-g14 | NA | (Vuong *et al.*, 2015) |
|  | 19 | 40218800 | Seed weight 9-g5.1 | NA | (Copley *et al.*, 2018) |
| SVR | 7 | 1032587 <sup>2</sup> | WUE 2-g18 | 2 | (Kaler *et al.*, 2017) |
|  | 19 | 40218800 | Seed weight 9-g5.1 | NA | (Copley *et al.*, 2018) |

<sup>a</sup> Detected in separate environments (1: 2018Ridgetown, 2:2019Ridgetwon, 3:2018Palmyra, 4:2019Palmyra, NA: Not found in any environment).
MLM: Mixed Linear Model, FarmCPU: Fixed and random model circulating probability unification, RF: Random Forest, SVR: Support Vector Regression

**Table 4.**
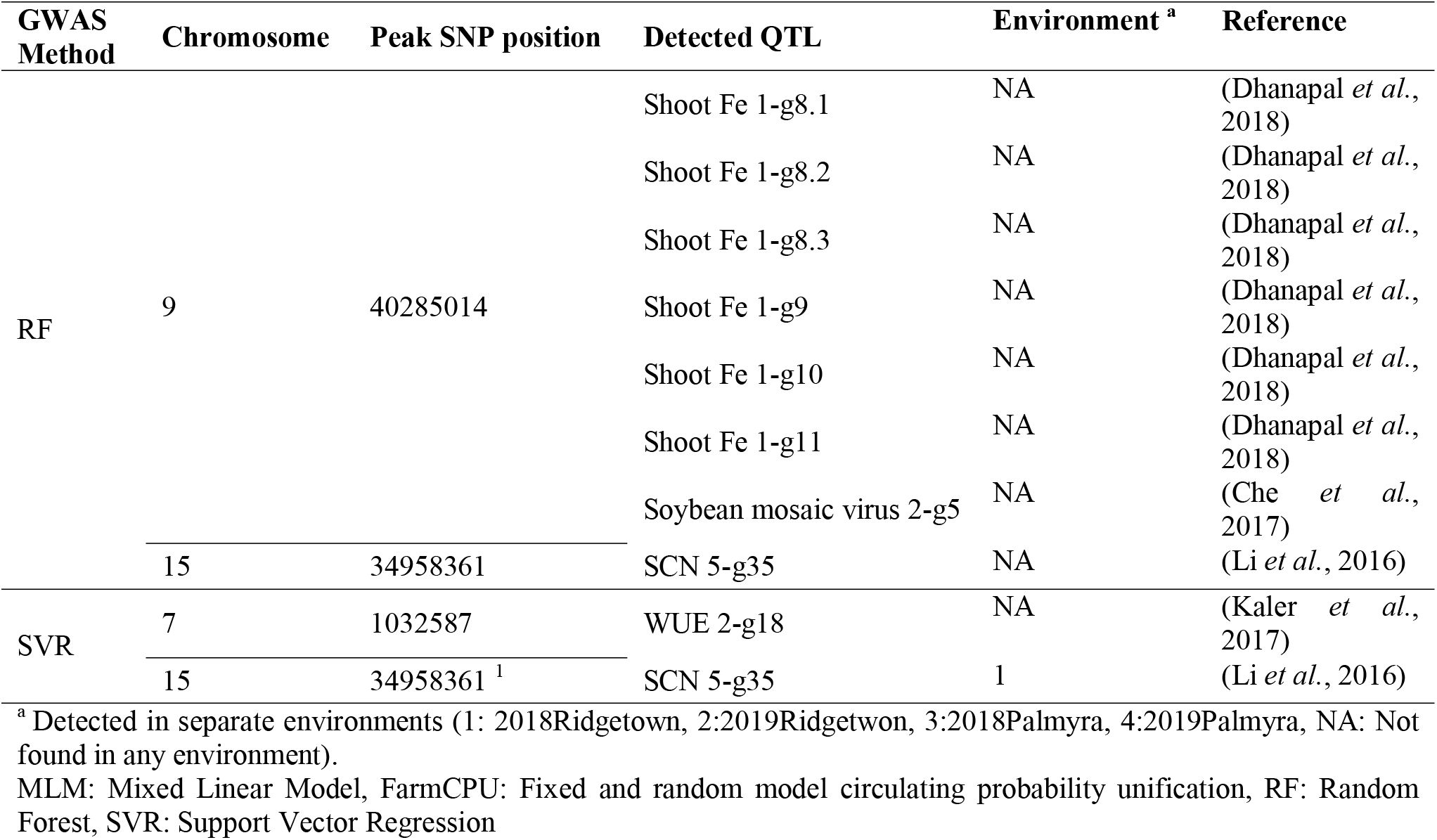
The list of detected QTL for soybean total number of reproductive nodes per plant (RNP) using different AS methods in the tested soybean population.

The average RNP was 11.89 nodes with a standard deviation of 0.98 nodes (Fig. 3). Based on the results of MLM and FarmCPU methods, four associated SNP markers with RNP were located on chromosomes 8 and 19. Using the RF method, four associated SNP markers were identified on chromosomes 8, 9, 15, and 20. Using the SVR method, 11 SNP markers were associated with RNP located on chromosomes 4, 7, 8, 15, 18, 19, and 20 (Fig. 7A). Regardless of the type of GWAS methods used in this study, we found SNP markers associated with the trait on chromosome 8. The position of the associated SNP marker on chromosome 8 was identical both in SVR and RF (462.3 Kbp) and MLM and FarmCPU (481.6 Kbp). The list of detected QTL for RNP is presented in Table 5. The average value for PP in the tested GWAS panel was 45.02 pods with a standard deviation of 8.54 pods. We did not detect any SNP marker associated with PP using the MLM and FarmCPU methods. However, by using the RF method, four SNP markers were found to be associated with PP and located on chromosomes 7, 10, 18, and 20 (Fig. 7B). Twelve associated SNP markers were found by SVR that were located on chromosomes 6, 9, 10, 11, 15, 18, and 19 (Fig. 7B). The GWAS of chromosome 10 with PP were found both in RF and SVR with 4.6 cM distance far from each other. In PP, MLM and FarmCPU did not detect any related QTL for this trait, while SVR-mediated GWAS was identified seven QTL directly related to the pod number (Table 6).

**Table 5.**
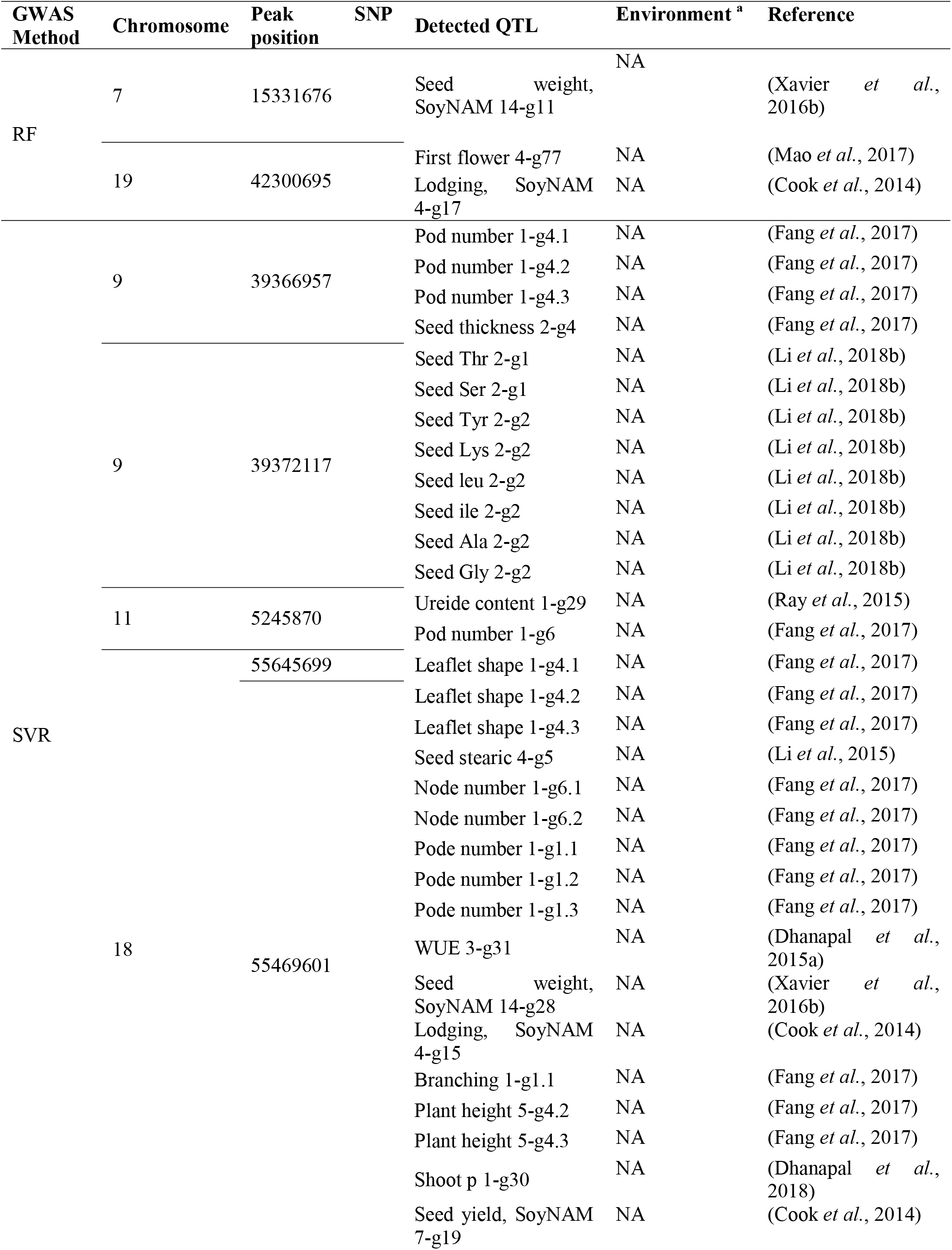
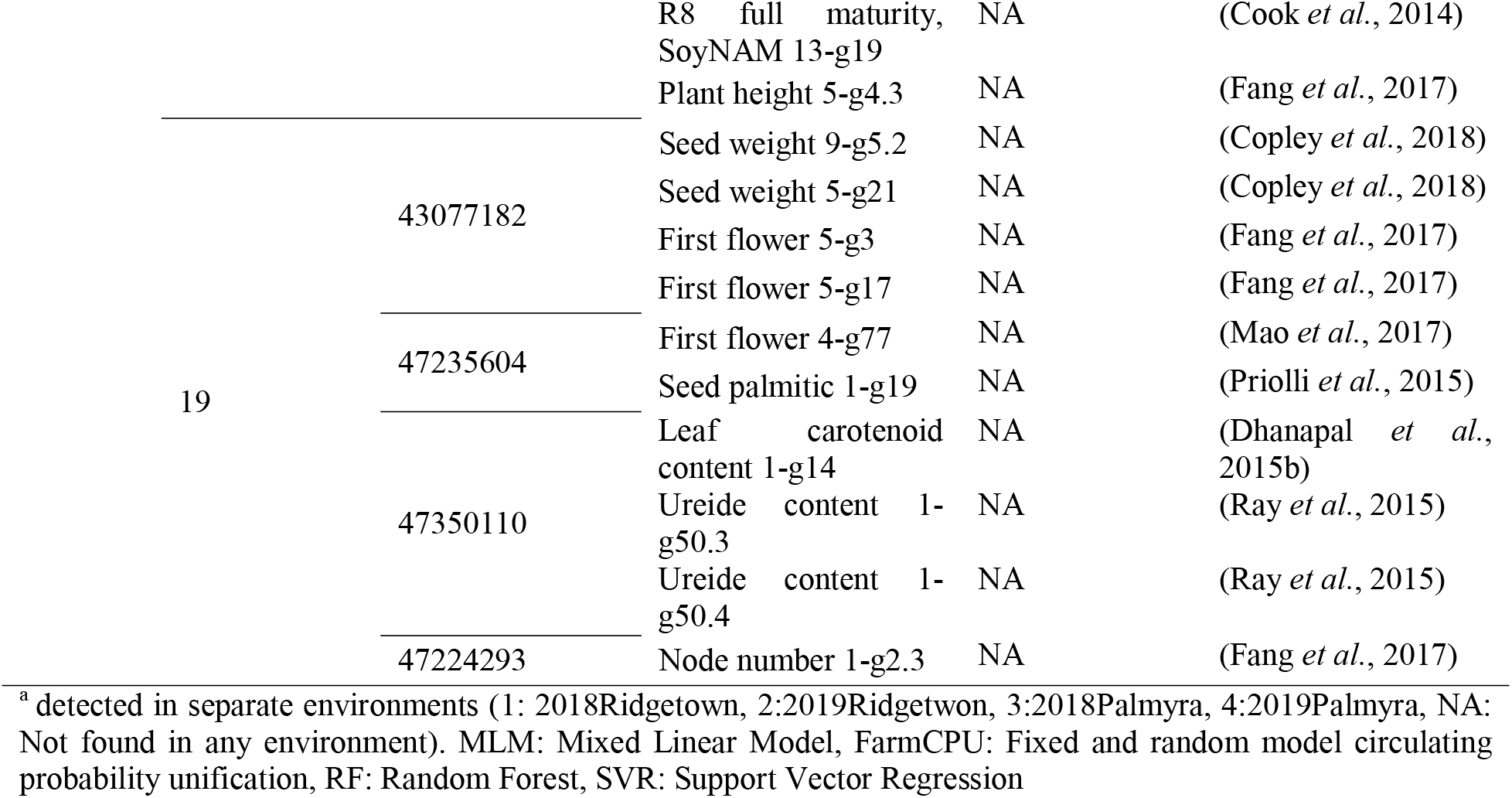
The list of detected QTL for soybean total number of pods per plant (PP) using different GWAS methods in the tested soybean population.

**Fig. 7.**
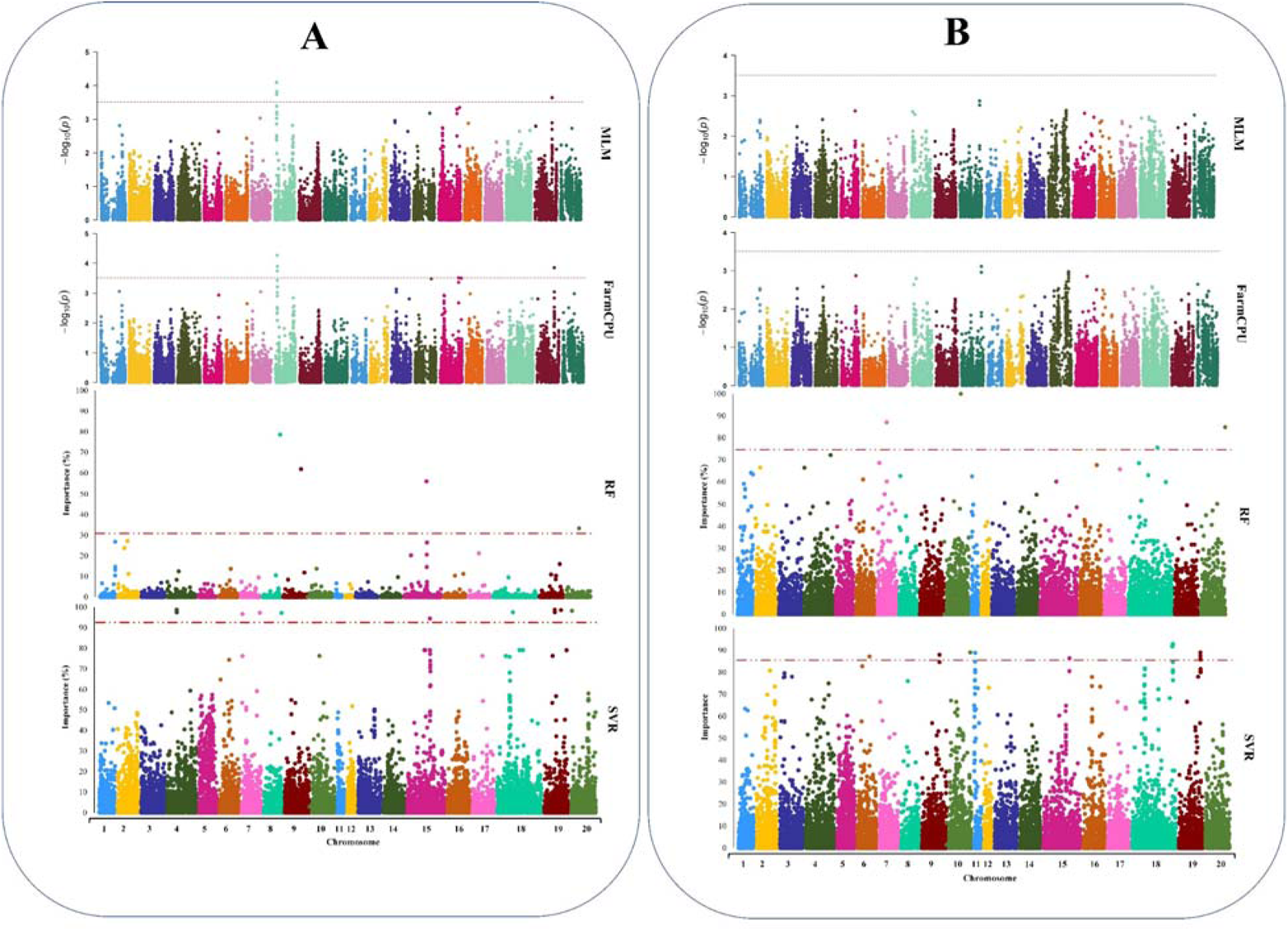
Genome-wide Manhattan plots for GWAS studies of A) The total number of reproductive nodes (RNP) and B) the total number of pods (PP) in soybean using MLM, FarmCPU, RF, and SVR methods, from top to bottom, respectively.

### Identification of candidate genes within QTL

According to the flanking regions of each QTL which was determined using LD decay distance, 150-kbp upstream and downstream of each SNP’s peak were considered to identify potential candidate genes (Fig. 1). Candidate genes were extracted for each significant peak SNP with high allelic effect and based on the gene annotation, enrichment tools and previous studies (Table S1). For maturity, three peak SNPs (Chr2_695362, Chr2_720134, and Chr19_47513536) had the highest allelic effect than other detected peak SNPs (Fig. 8A). On the basis of the gene annotation and expression within QTL, *Glyma.02g006500* (GO:0015996) and *Glyma.19g224200* (GO:0010201) were identified as the strong candidate genes for maturity, which encode chlorophyll catabolic process and phytochrome A (PHYA) related genes, respectively. *Glyma.02g006500* (GO:0015996) was exactly detected in the peak SNP position of Chr2_695362, whereas *Glyma.19g224200* (GO:0010201) was 119 Kbp far from the detected peak SNP at Chr19_47513536. In yield, the peak SNP with the position of Chr7_1032587 had the highest allelic effect in comparison with other detected peak SNPs (Fig. 8B). Within a 77 Kbp above from the detected peak SNP (Chr7_1032587), *Glyma.07G014100* (GO:0010817) was identified, which encodes the regulation of hormone levels, as the strongest candidate genes in yield. For NP, two peak SNPs (Chr7_1032587 and Chr7_1092403) had the highest allelic effect among all detected peak SNPs (Fig. 8C). SNP peak position of Chr7_1032587 was detected in common for yield, NP, and NRNP. *Glyma.07G205500* (GO:0009693) and *Glyma.08G065300* (GO:0042546) were detected as the strongest candidate genes both in NP and NRNP, which encode UBP1-associated protein 2C and cell wall biogenesis, respectively. Both detected gene candidates were exactly at the associated peak SNPs at Chr7_1032587 and Chr8_5005929 (Fig. 8D). Regarding peak SNPs associated with RNP, the highest allelic effects were found in peak SNPs of Chr9_40285014 and Chr15_34958361 (Fig. 8E). *Glyma.15G214600* (GO:0009920) and *Glyma.15G214700* (GO:0009910), which encode cell plate formation involved in plant-type cell wall biogenesis and acetyl-CoA biosynthetic process, as strong candidate genes in NRNP. *Glyma.15G214600* (GO:0009920) and *Glyma.15G214700* (GO:0009910) were 127 and 90 Kbp far from the detected peak SNP at Chr15_3495836, respectively. In PP, the highest allelic effects were found in peak SNPs at Chr7_15331676, Chr11_5245870, and Chr18_55469601 (Fig. 8F). *Glyma.07G128100* (GO:0009909) was the strongest candidate genes for PP, which encodes regulation of flower development. *Glyma.07G128100* (GO:0009909) was detected exactly in the peak SNP position of Chr7_15331676.

**Fig. 8.**
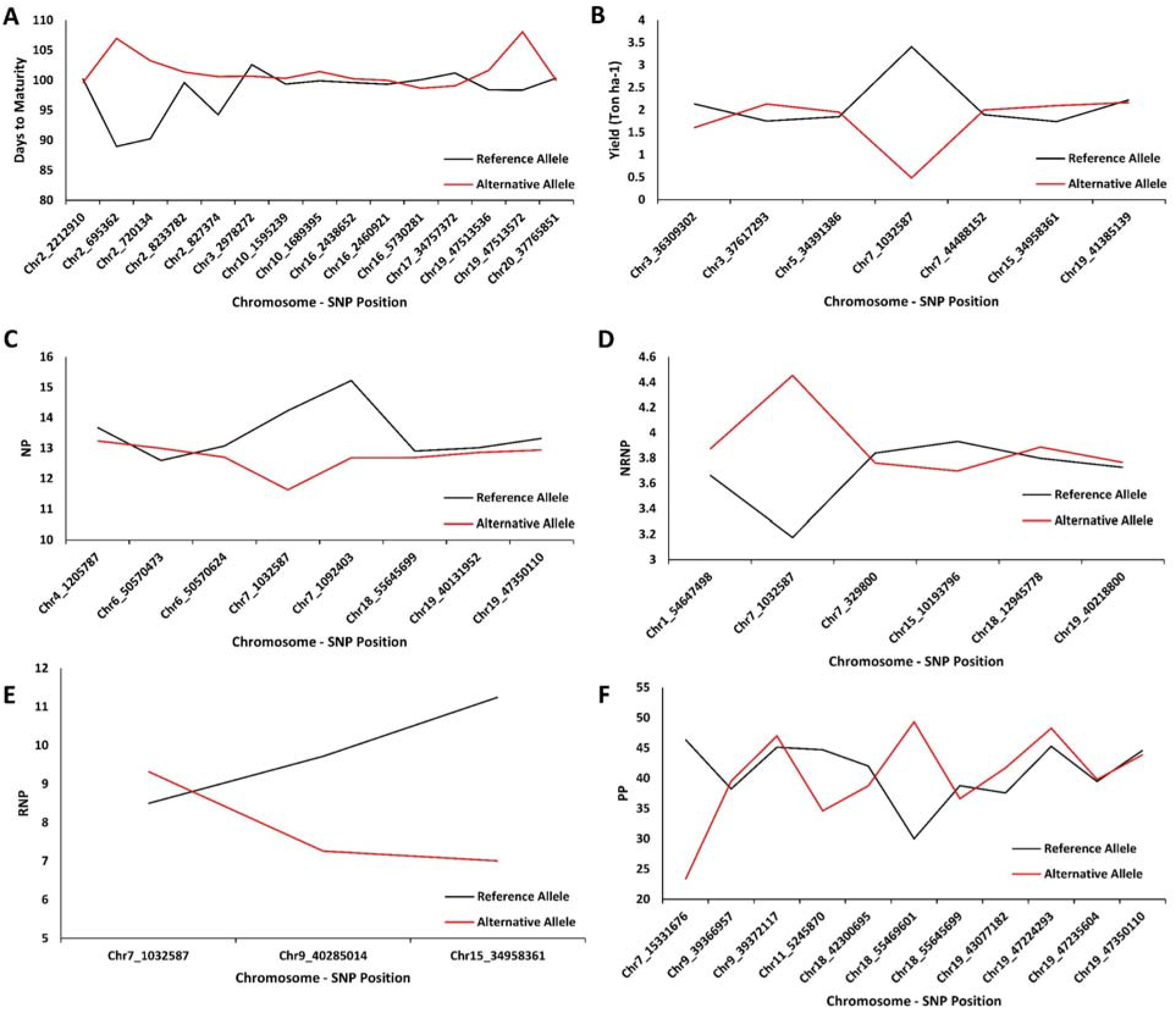
The average effects of reference allele and alternative allele from the detected SNP’s peak for seed yield (A), maturity (B), NP (C), NRNP (D), RNP (E), and PP (F) in 227 soybean genotypes across four environments. RNP: Total number of reproductive nodes per plant, NRNP: The total number of non-reproductive nodes per plant, NP: The total nodes per plant, PP: The total number of pods per plant

## Discussion

One of the objectives of this study was to gain a better understanding of the roles of soybean yield component traits in the production of total seed yield and how these traits can be used for facilitating the development of high-yielding soybeans. The genetic dissection of soybean yield component traits in order to develop genetic and genomics toolkits can be useful for designing breeding population and selection criteria aiming at improving yield genetic gains in new cultivars (Cooper *et al.*, 2009; Hu *et al.*, 2020; Xavier and Rainey, 2020). For this aim, a wide range of analyses, including Pearson correlation, normality and distribution plots, GWAS both in combined and separate environments, and functional annotation of candidate genes and QTL, were performed in this study. The collective evaluation of the mentioned analysis contributes to building the wide perspectives of the genetic architecture of the soybean yield component traits. One of the important factors for genetic studies is to evaluate the phenotypic variation within genotypes and environments. High phenotypic variation was observed for yield and PP, while maturity and NP had the lowest phenotypic variation across the tested environments. These findings are in line with the results of previous research on yield component traits (Kahlon and Board, 2012; Xavier and Rainey, 2020). The heritability and correlation analyses showed that NP had the highest heritability and significant linear correlations with RNP and PP. Also, PP had the highest correlation with yield among all the tested soybean yield components. The number of nodes and pods in soybean are known as the two of the key soybean yield components that play an important role in determining the final soybean seed yield (Herbert and Litchfield, 1982; Kahlon and Board, 2012; Xavier and Rainey, 2020). However, studies showed the low heritability rates for soybean yield components, especially NP and PP (KUSWANTORO, 2017; Sulistyo and Sari, 2018; Xavier *et al.*, 2016a; Xavier and Rainey, 2020). The nature of these traits can explain low heritability rates as they are mostly affected by environmental factors (Price and Schluter, 1991). Although heritability indicates the strength of the relationship between phenotype and genetic variation of the particular trait, it does not indicate the value of the trait for genetic study (Cassell, 2009). Different low heritable traits are highly correlated with significant economic traits (Cassell, 2009). In soybean, yield can be considered as the most important economic trait that is highly determined by yield components. Therefore, any genetic and environmental studies around yield components can open the possibility of overall yield improvement in major crops such as soybean.

GWAS is known as one of the most important genetic toolkits for detecting QTL associated with quantitate traits (Kaler *et al.*, 2020). There are several statistical methods implemented in GWAS for improving the detection of associated SNP markers with the trait of interest. While conventional GWAS are appropriate approaches for detecting SNP markers with large effects on complex traits, they are, however, underpowered for the simultaneous consideration of a wide range of interconnected biological processes and mechanisms that shape the phenotype of complex traits (Lee *et al.*, 2020). Therefore, using variable importance values in ML algorithms for identifying SNP-trait associations may improve the power of ML-mediated GWAS for discovering variant-trait association with higher resolution (Szymczak *et al.*, 2009). The variable importance methods based on linear and logistic regressions, support vector machines, and random forests are well established in the literature (Grömping, 2009; Williamson *et al.*, 2020; Wu and Liu, 2009; Yoosefzadeh-Najafabadi *et al.*, 2021a). Among all the tested GWAS methods in this study, SVR-mediated GWAS was the best method to detect SNP markers with high allelic effects associated with the tested traits. The advantage of SVR-mediated GWAS over conventional GWAS models can be explained by the presence of a nonlinear relationship between input and output variables, which is used to build an algorithm with accurate prediction ability (Kaneko, 2020). Therefore, genomic regions could be better detected by SVR-mediated GWAS because of its ability to consider the interaction effects between SNPs rather than *p*-values for individual SNP-trait GWAS tests.

None of the detected QTL by MLM, FarmCPU, and RF were reported to be associated directly with soybean maturity. However, using SVR-mediated GWAS, five QTL were detected on chromosomes 16 and 19 specifically related to the soybean maturity. Those QTL were previously reported by Sonah *et al.* (2015) and Copley *et al.* (2018) in separate studies. Also, the peak SNP position of Chr19_47513536 detected by SVR-mediated GWAS had the highest allelic effect among all the detected SNPs in soybean maturity, which is in line with Sonah *et al.* (2015). For soybean seed yield, five QTL detected by SVR-mediated GWAS were reported previously (Copley *et al.*, 2018; Hu *et al.*, 2014), while none of the detected QTL from other tested GWAS methods was previously reported for this trait. There was no previous study on the genetic structure of NRNP and RNP, therefore, all the detected QTL in this study are presented for the first time. For PP, conventional GWAS methods were not able to detect any associated QTL. However, using SVR-mediated GWAS, a total of seven QTL were detected to be related to pod numbers based on previous studies (Zhang *et al.*, 2015a). It would be necessary to emphasize that the average allelic effects of all detected QTL presented in Fig. 8 was not directly estimated by the tested GWAS methods. The RF and SVR-mediated GWAS methods do not specifically provide an allele effect therefore, the aim of this study was mostly focused on detecting the associated genes and QTL underlying the soybean yield, maturity, and yield components.

The results of candidate gene identifications within identified QTL by SVR-mediated GWAS analyses reveled important information. For example, from all the detected genes using SVR-mediated GWAS for maturity, candidate gene *Glyma.02g006500* (GO:0015996) is a protein ABC transporter 1, that is annotated as a chlorophyll catabolic process and located exactly in the peak SNP position at Chr02_695362. ATP-binding cassette (ABC) transporter genes play conspicuous roles in different plant growth and developmental stages by transporting different phytochemicals across endoplasmic reticulum (ER) membranes (Hwang *et al.*, 2016). Because of the central roles of ABC transporters in transporting biomolecules such as phytohormones, metabolites, and lipids, they play important roles in plant growth and development as well as maturity (Block and Jouhet, 2015; Hwang *et al.*, 2016). Moreover, recent studies revealed that ER uses fatty acid building blocks made in the chloroplast to synthesize Triacylglycerol (TAG). Therefore, ABC transporter genes are important for the normal accumulation of Triacylglycerol (TAG) during the seed-filling stage and maturity (Block and Jouhet, 2015; Kim *et al.*, 2013). Additionally, *Glyma.19g224200* (GO:0010201) in E3 locus, which was previously discovered by Buzzell (1971) and molecularly characterized as a phytochrome A (PHYA) gene (Watanabe *et al.*, 2009), was detected through the SVR-mediated GWAS. Phytochromes, through PHYTOCHROME INTERACTING FACTOR (PIF), regulate the expression of some specific genes encoding rate-limiting catalytic enzymes of different plant growth regulators (e.g., abscisic acid, gibberellins, auxin) and, therefore, play crucial roles in plant maturity (Legris *et al.*, 2019). In addition, PHYB is inactivated after imbibition shade signals, which repress PHYA-dependent signaling in the embryo that results in the maturity of seeds by preventing germination (Casal, 2013; De Wit *et al.*, 2016). This is obtained by regulating the balance between abscisic acid and gibberellin. Subsequently, abscisic acid transports from the endosperm to the embryo by ABC transporter (De Wit *et al.*, 2016).

Regarding NRNP, candidate gene *Glyma.07G205500* (GO:0009693-UBP1-associated protein 2C) that annotated as ethylene biosynthetic process was located exactly in the peak SNP position of *Chr7_37469678*, was detected by SVR-mediated GWAS. An interaction screen with the heterogeneous nuclear ribonucleoprotein (hnRNP) results in the production of oligouridylatebinding protein 1 (UBP1)-associated protein (Lambermon *et al.*, 2002). It has been well documented that this protein plays important roles in several physiological processes such as responses to abiotic stresses (Li *et al.*, 2002), leaf senescence(Kim *et al.*, 2008), floral development (Streitner *et al.*, 2008), and chromatin modification (Liu *et al.*, 2007). In addition, previous studies showed that the production of productive or non-reproductive nodes is completely accompanied by the upregulation or downregulation of this protein (Bäurle and Dean, 2008; Na *et al.*, 2015). In addition, *Glyma.08G065300* (GO:0042546-MADS-box transcription factor) that is associated with cell wall biogenesis, was located in the SNP position of Chr8_5005929. The genes of the MADS-box family can be considered as the main regulators for cell differentiation and organ determination (Lee *et al.*, 2013). The floral organ recognition MADS-box family has been categorized into A, B, C, D, and E classes. Among these classes, class E was shown to be associated with reproductive organ development (Hussin *et al.*, 2021). Indeed, activation or repression of this transcription factor leads to the development of nodes to productive or non-productive nodes (Ditta *et al.*, 2004; Gao *et al.*, 2010; Liu *et al.*, 2013).

Gene expression data provided by Severin *et al.* (2010) noted that 20 candidate genes for PP that were detected using the SVR-mediated GWAS were expressed in flowers, 1 cm pod (7 DAF), pod shell (10-13 DAF), pod shell (14-17 DAF) and seeds. In PP, most of the genes detected by SVR-mediated GWAS are associated with auxin influx carrier or auxin response factors (ARFs), gibberellin synthesis, and response to brassinosteroid (Lin *et al.*, 2020; Yin *et al.*, 2018). Song *et al.* (2020) and Li *et al.* (2018a) also reported that some genes related to PP were associated with embryo development, stamen development, ovule development, cytokinin biosynthesis, and response gibberellin that we also identified in our study. Soybean seed yield significantly depends on seed number and seed size (Liu *et al.*, 2010; Rotundo *et al.*, 2009). These two factors are determined from fertilization to seed maturity. Therefore, soybean seed development can be divided into three stages or phases: pre-embryo or seed set, embryo growth or seed growth, and desiccation stages or seed maturation phases (Ruan *et al.*, 2012; Weber *et al.*, 2005). In Arabidopsis, a complex signaling pathway and regulatory networks, including sugar and hormonal signaling, transcription factors, and metabolic pathway, have been reported to be involved in seed development (Le *et al.*, 2010; Orozco-Arroyo *et al.*, 2015). Several key genes and transcription factors (e.g., LEAFY COTYLEDON 1 (LEC1), LEC2, FUSCA3 (FUS3), AGAMOUS-LIKE15 (AGL15), ABSCISIC ACID INSENSITIVE 3 (ABI3), YUCCA10 (YUC10), ARFs) have been determined to control several downstream plant growth regulators pathways to the seed development (Lepiniec *et al.*, 2018; Pelletier *et al.*, 2017; Sun *et al.*, 2010). Indeed, a high ratio of abscisic acid to gibberellic acid can regulate seed development (Figueiredo and Köhler, 2018; Wang *et al.*, 2016). The downregulation of FUS3 obtains this through repressing GA3ox1 and GA3ox2 and activating ABA biosynthesis (Weber *et al.*, 2005). In soybean, RNA seq analysis for the seed set, embryo growth, and early maturation stages of developing seeds in two soybeans with contrasting seed size showed cell division and growth genes, hormone regulation, transcription factors, and metabolic pathway are involved in seed size and numbers (Du *et al.*, 2017).

## Conclusion

A better understanding of the genetic architecture of the yield component traits in soybean may enable breeders to establish more efficient selection strategies for developing high-yielding cultivars with improved genetic gains. Major yield components such as maturity, NP, NRNP, RNP, and PP play important roles in determining the overall yield production in soybean. This study verified the importance of those traits, using correlation and distribution analyses, in determining of the total soybean seed yield. Furthermore, by testing different conventional and ML-mediated GWAS methods, this study demonstrated the potential benefit of using ML-mediated methods in GWAS. SVR-mediated GWAS outperformed all the other methods tested in this study, and therefore, it is recommended as an alternative to conventional GWAS methods with a greater power for detecting genomic regions associated with complex traits such as yield and its components in soybean, and possibly other crop species. To the best of our knowledge, this study is the first attempt in which SVR was used for GWAS analyses in plants. In order to verify the causal relationship between identified QTL and the target phenotypic traits, we identified candidate genes within each QTL using gene annotation procedures and information. The results demonstrated the efficiency of SVR-mediated GWAS in detecting reliable QTL that can be used in marker-assisted selection. Nevertheless, further investigation is recommended to confirm the efficiency of SVR-mediated GWAS in detecting associated genomic regions in other plant species.

## Supplementary Data

**Table S1** The full list of detected genes using different GWAS methods for soybean seed yield, maturity, and yield component traits.

## Acknowledgments

The authors are grateful to the past and current members of Eskandari laboratory at the University of Guelph, Ridgetown, Bryan Stirling, John Kobler, and Robert Brandt for their technical support. We would like to thank Maryam Vazin and Mohsen Hesami for their assistance with the field data collection and reviewing the manuscript, respectively.

## Author Contribution

ME conceptualized, designed and directed the experiments. MY-N performed the experiments, modeled, summed up, and wrote the manuscript. ST participated in candidate gene analyses; DT, DTOR, ST, IR, and ME revised the manuscript and validated the results. All authors have read and approved the final manuscript.

## Data Availability Statement

The raw data supporting the conclusions of this article will be made available by the authors, without undue reservation.

